# Caterpillars Count! A citizen science project for monitoring foliage arthropod abundance and phenology

**DOI:** 10.1101/257675

## Abstract

Caterpillars Count! is a citizen science project that allows participants to collect data on the seasonal timing, or phenology, of foliage arthropods that are important food resources for forest birds. This project has the potential to address questions about the impacts of climate change on birds over biogeographic scales. Here, we provide a description of the project’s two survey protocols, evaluate the impact of survey methodology on results, compare findings made by citizen scientist participants versus trained scientists, and identify the minimum levels of sampling frequency and intensity in order to accurately capture phenological dynamics. We find that beat sheet surveys and visual surveys yield similar relative and absolute density estimates of different arthropod groups, with beat sheet surveys recording a higher frequency of beetles and visual surveys recording a higher frequency of flies. Citizen scientists generated density estimates within 6% of estimates obtained by trained scientists regardless of survey method. However, patterns of phenology were more consistent between citizen scientists and trained scientists when using beat sheet surveys than visual surveys. By subsampling our survey data, we found that conducting 30 foliage surveys on a weekly basis led to 95% of peak caterpillar date estimates to fall within one week of the "true" peak. We demonstrate the utility of Caterpillars Count! for generating a valuable dataset for ecological research, and call for future studies to evaluate how training and resource materials impact data quality and participant learning gains.

---

One of the observed impacts of climate change over recent decades has been a shift in the seasonal timing, or phenology, of organisms and their life cycles. For example, first flowering dates in Concord, Massachusetts have advanced by two to three weeks since Thoreau’s records from the 1850s (Ellwood et al., 2013; Primack, 2014). Butterflies have similarly advanced first flight dates over recent decades (Altermatt, 2012; Forister and Shapiro, 2003), and many bird species have advanced the timing of migration (Hurlbert and Liang, 2012; Mayor et al., 2017). Such observed phenological shifts indicate that these species are able to respond to changes in their physical environment, and yet the magnitude of these shifts is highly variable among species and across trophic levels (Both et al., 2009; Parmesan, 2007; Parmesan and Yohe, 2003). Phenological mismatch occurs when organisms fail to adjust phenologically to the same degree as the organisms on which they depend, and has been documented between plants and their pollinators (Forrest, 2015), insects and their host plants (Singer and Parmesan, 2010), and birds and the arthropods they rely on for successfully raising offspring (Visser et al., 2006, 2012). Understanding phenological mismatch in migratory birds is a particularly challenging problem because these birds often traverse thousands of kilometers, and climate change is geographically variable over these regions. As such, observed phenological shifts in the northeastern US, for example, may have little correlation with phenological shifts in the southeast, and yet whether these changes are correlated may have important impacts on migratory birds (Fontaine et al., 2015; Wood and Kellermann, 2015).

Citizen science programs are one of the most effective ways to monitor simple biological phenomena like phenology over broad geographic extents as demonstrated by the recent efforts by the National Phenology Network (Schwartz et al., 2012), Project Budburst (Johnson, 2016), and eBird (Sullivan et al., 2014). Individual scientists or research groups are simply unable to collect data efficiently at the relevant spatial and temporal scales for addressing these broad biogeographical questions. Here we introduce a new citizen science project, *Caterpillars Count!* (http://caterpillarscount.unc.edu), whose aim is to document geographic and annual variation in the phenology and abundance of arthropods that foliage gleaning birds rely on during the breeding season. The name of the project highlights the fact that Lepidoptera larvae in particular represent an important and often primary food source (Holmes et al., 1979; Holmes and Schultz, 1988; Jones et al., 2003; Sillett et al., 2000) known to influence avian density (Graber and Graber, 1983), reproductive success (Rodenhouse and Holmes, 1992; Visser et al., 2006), clutch size (Perrins, 1991) and number of broods raised (Nagy and Holmes, 2005a, 2005b). The enlistment of citizen scientists would potentially allow for an examination of phenological mismatch between birds and their food resources at an unprecedented scale.

Our aims in this paper are to 1) describe the survey protocols used to monitor foliage arthropods, 2) evaluate the impact of survey methodology on results, 3) compare findings made by citizen scientist participants versus trained scientists to assess the reliability of citizen science data collection and to make recommendations for citizen science coordinators, and 4) identify the minimum levels of sampling frequency and intensity in order to accurately capture phenological dynamics. It is our hope that *Caterpillars Count!* will yield robust data on arthropod phenology over broad spatial scales that can ultimately be leveraged with other existing datasets to provide new insights into potential mismatches between vegetation, arthropods, and birds.

## *Caterpillars Count!* Protocol

Because arthropods may be patchily distributed across an area, accurate estimates of density require conducting many surveys per survey date. Permanent survey locations are arrayed across the study site in groups (“circles”) of five, with a central survey branch identified opportunistically (e.g., a branch with lots of additional suitable vegetation nearby) followed ideally by the first suitable branch 5 m away in each of the four cardinal directions (Figure 1, inset). To be suitable, a branch must have at least 50 leaves (or leaflets for compound leaves) each greater than 5 cm in length. Participating sites may have anywhere from 20 to 60 surveys arranged in 4 to 12 circles across the study site.

**Figure 1:**
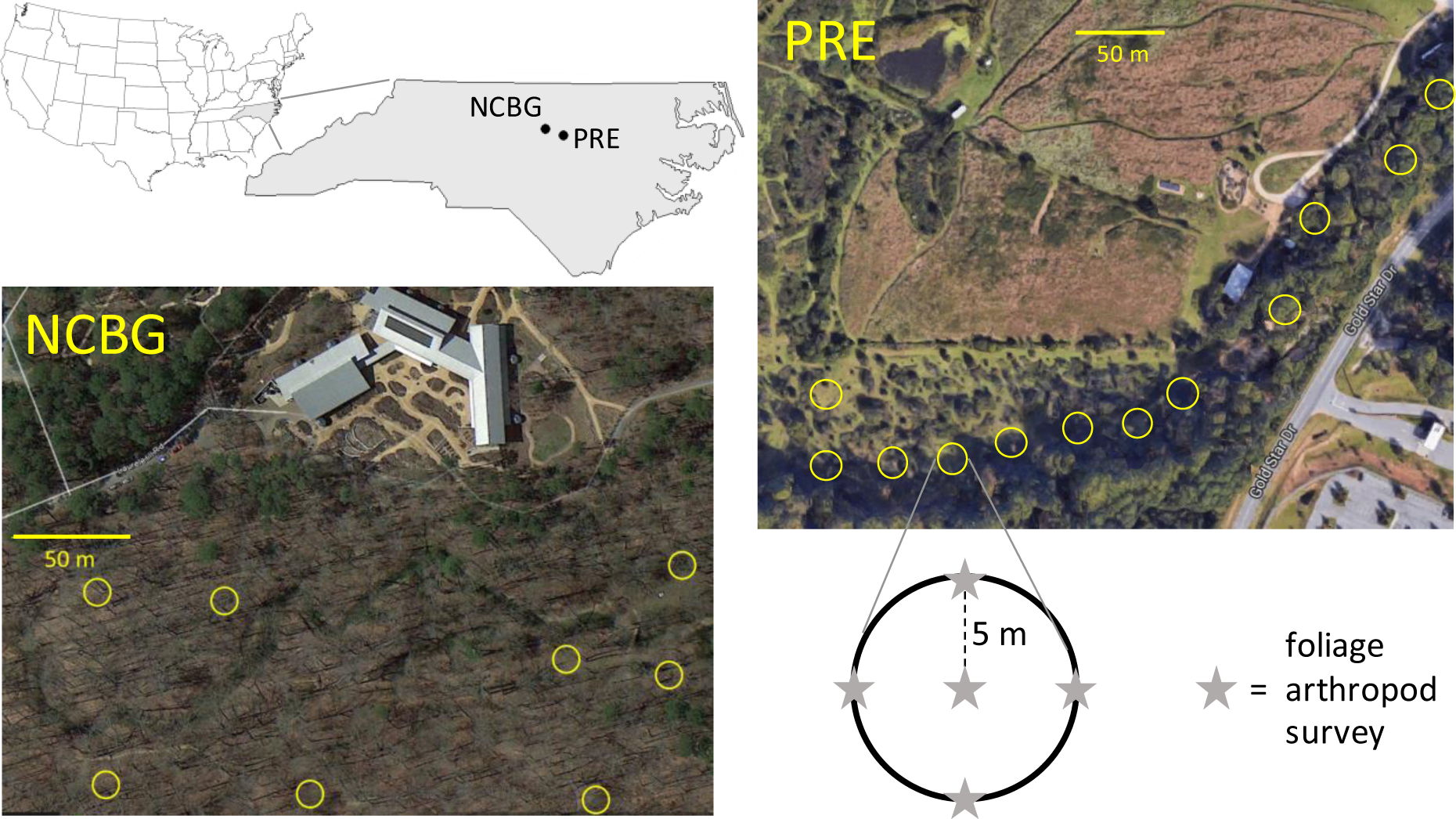
Location of North Carolina Botanical Garden (NCBG) and Prairie Ridge Ecostation (PRE) within North Carolina, and the layout of survey circles at each site. Each survey circle consists of 5 foliage arthropod surveys. Photos from Google Earth.

### Visual foliage survey

Visual foliage surveys conducted at ground level have been used for decades to characterize foliage arthropod availability to birds throughout the forest canopy (Holmes and Schultz, 1988).

For one survey, an observer examines both the upper- and undersides of 50 leaves and associated petioles and twigs on a branch of woody vegetation typically 1-2 m above the ground. All arthropods observed greater than 2 mm in length are identified, generally to order (but in some cases suborder or family; Table 1), and their body length (not including legs or antennae) is recorded to the nearest millimeter. Arthropods smaller than 2 mm are ignored both because of their lesser importance as food items as well as the increased difficulty and therefore time required for identification. A single visual foliage survey takes 2-6 minutes depending upon the density of arthropods, experience of the observer, and degree of clustering of leaves on a branch.

**Table 1.** Common arthropod groups found on foliage that citizen scientist participants are expected to be able to identify.

| Common name | Scientific name | Taxonomic level | Distinguishing features |
| --- | --- | --- | --- |
| Ants | Formicidae | Family | Narrow waist, no wings; elbowed antennae |
| Aphids and psyllids | Sternorrhyncha | Suborder, Order Hemiptera | Small (just a few mm); aphids are pear-shaped |
| Bees and wasps | Hymenoptera (excluding Formicidae) | Order | 2 pairs of wings with the hindwings smaller than the frontwings; wasps have narrow waists but bees do not |
| Beetles | Coleoptera | Order | A straight line down the back where the two hard wing casings (elytra) meet |
| Caterpillars | Lepidoptera (larvae) | Order | Soft, cylindrical body with 6 legs and up to 5 pairs of prolegs |
| Daddy longlegs | Opiliones | Order | 8 very long legs; they appear to have a single oval-shaped body |
| Flies | Diptera | Order | A single pair of wings |
| Grasshoppers, Crickets | Orthoptera | Order | Usually with enlarged hind legs for jumping |
| Leafhoppers, Cicadas | Auchenorrhyncha | Suborder, Order Hemiptera | Usually a wide head relative to the body; hoppers have wings folded tentlike over their back, while cicadas have large membranous wings. |
| Moths, Butterflies | Lepidoptera (adults) | Order | 4 large wings covered by fine scales |
| Spiders | Araneae | Order | 8 legs, with two distinct body segments: the cephalothorax and abdomen |
| True Bugs | Heteroptera | Suborder, Order Hemiptera | Semi-transparent wings which partially overlap creating a triangle or X shape on the back; often has pointy "shoulders" |

### Beat sheet survey

As an alternative to the visual foliage survey, participants may choose instead to conduct a beat sheet survey in which the survey branch is beat with a stick ten times in rapid succession over a white 60 x 60 cm sheet. As with the visual survey, all arthropods are identified to the relevant order/group (Table 1) and length is recorded to the nearest millimeter. In addition, the participant records the total number of leaves that were positioned above the beat sheet during beating which is expected to vary from branch to branch. A single beat sheet survey typically takes 2-3 minutes depending on the density of arthropods and experience of observer.

## Methods

### Data collection

Foliage arthropod surveys were conducted at two locations within the North Carolina Piedmont region. The North Carolina Botanical Garden site (NCBG; 35.898550° N, 79.031642° W) is a natural deciduous forest in Chapel Hill, NC featuring a canopy dominated by *Fagus grandifolia* and *Acer sacharrum* with an understory of *Lindera benzoin* and *Carpinus caroliniana.* Prairie Ridge Ecostation (PRE; 35.8117° N, 78.7139° W) is an outdoor nature center in Raleigh, NC featuring a narrow forest strip (including *Liquidambar styraciflua, Acer negundo, Diospyros virginiana)* alongside an open prairie. Forty survey locations were established at NCBG and 60 at PRE (Figure 1).

In both 2015 and 2016, members of the Hurlbert Lab at the University of North Carolina (hereafter “trained scientists”) conducted visual and beat sheet surveys twice per week from mid-May through July at all survey locations within NCBG and PRE. Hurlbert provided extensive training before and during foliage survey activities, ensuring that team members were capable of documenting potentially cryptic arthropods and of identifying arthropods to the relevant groups. Visual surveys were conducted first at each survey location followed by a beat sheet survey on an adjacent branch of the same plant species. Surveys were typically conducted between 0830 and 1200 hrs. At PRE in 2015, trained scientists additionally conducted beat sheet surveys once per week on Thursday afternoons, typically between 1300 and 1400, at a fixed subset of 40 of the 60 total survey locations.

In both 2015 and 2016, volunteers (hereafter “citizen scientists”) were recruited to conduct foliage arthropod surveys at PRE at the fixed subset of 40 survey locations. Citizen scientists were recruited through the volunteer program at the North Carolina Museum of Natural Sciences and included both men and women varying in age from 22 to 50 years in age. Volunteers were trained by CLG, who worked with the volunteers the first three times they conducted surveys and focused heavily on arthropod identification skill building. After the third survey, the volunteers conducted the surveys on their own. In 2015, seven different citizen scientists conducted visual foliage surveys, some on Thursdays between 1300 and 1500 and others on Saturdays between 0900 and 1100 hours most weeks. In 2016, four citizen scientists were recruited, and conducted beat sheet surveys once per week on average, typically between 0800 and 1200 hrs. We were thus able to compare citizen scientist and trained scientist observations based on visual surveys in 2015 and based on beat sheet surveys in 2016. An average citizen scientist conducted 140-280 surveys over the course of each season, while each trained scientist typically conducted 900-1400 surveys per season and so had more experience on top of the increased training and supervision.

Finally, while our survey methodology focuses on foliage 1-2 m above ground for logistical reasons, we would ideally like to make inferences about arthropod phenology throughout the entire canopy. In order to validate this comparison between foliage strata, we collected caterpillar frass falling from the canopy at both sites in 2015 to compare with observed phenology from the ground level foliage surveys. Frass traps consisted of a 20 cm diameter plastic funnel mounted onto a garden stake 30 cm above ground level and lined with a 40 cm diameter piece of filter paper folded into a cone. Each frass trap samples a cross-sectional area of 1662 cm^2^. Frass traps were located within existing survey circles (1 trap per circle at PRE, 2 per circle at NCBG) such that they spanned the same locations as the arthropod surveys. Although frass traps were collected and reset every 3-4 days, data were unusable on dates where there had been major rainstorms since the traps were deployed.

### Data analysis

Although participants recorded observations of all arthropods at least 2 mm in length, we only used observations of arthropods 5 mm long or longer in analyses. This reduces the incidence of misidentification of very small individuals, and also minimizes the effect of error in estimating the 2 mm cutoff. Using visual foliage survey data from trained scientists we calculated the average density per 50-leaf survey of each arthropod group by tree species.

Comparisons of relative arthropod composition between survey methods and between survey participant groups was conducted using chi-squared analyses, while comparisons of absolute density (number observed per survey) across all arthropod groups were conducted using Pearson’s correlation coefficients.

Phenology was characterized by the fraction of surveys (occurrence) on which a focal arthropod group was detected on a given date. We used occurrence rather than mean density estimates because the latter are sensitive to outliers, and we had a few instances in which a large number of gregarious caterpillars were observed in a single survey. Because citizen scientists typically collected data only once per week, we averaged the bi-weekly samples of trained scientists into weekly estimates in order to visually compare phenology and calculate Pearson’s correlation coefficients across weeks.

In order to assess the impact of sampling intensity and sampling frequency on estimates of peak caterpillar phenology date, we used data from Prairie Ridge in 2015 where trained scientists conducted 60 beat sheet surveys twice per week from mid-May through mid-July. We fit a Gaussian curve to these data (excluding the last two dates in July which reflect a late season peak less relevant for the avian breeding season; see Figure 3a below) and assumed the estimated mean of this curve reflected the “true” peak date (julian day 172). We then randomly subsampled the full dataset by manipulating both the number of surveys examined per sampling date (10, 20, 30, 40, 50, or 60 out of the 60 surveys) and the sampling frequency (every sampling date used, every other, every third, every fourth, and every fifth). For each combination of survey number and sampling frequency we conducted 60 replicate subsamples evenly split across potential starting dates (i.e., if sampling frequency was set at every other sampling date, we subsampled using the 1st, 3rd, 5th, etc. dates, but as another replicate also the 2nd, 4th, 6th, etc.). We estimated the peak date from Gaussian fits to the subsampled data. Fits were only used if the mean date was between julian days 100 and 200, and if the R^2^ for the fit was >0.2 (89% of all fits).

## Results

### Relative and absolute density

Visual foliage arthropod surveys revealed differences in the total density and composition of arthropods supported by different tree species (Figure 2a). Of the tree species with at least 300 surveys each, sweetgum (*Liquidambar styraciflua*) supported the highest arthropod density while sugar maple *(Acer saccharum)* had the lowest. Sweetgum and American beech *(Fagus grandifolia)* supported the greatest densities of caterpillars (0.14 and 0.17 caterpillars per survey, respectively, excluding one sweetgum survey with a colony of 250 fall webworm *(Hyphantria cunea)* caterpillars), although nearly all of the caterpillars on American beech were hidden between two leaves sewn together and so potentially inaccessible to birds.

**Figure 2:**
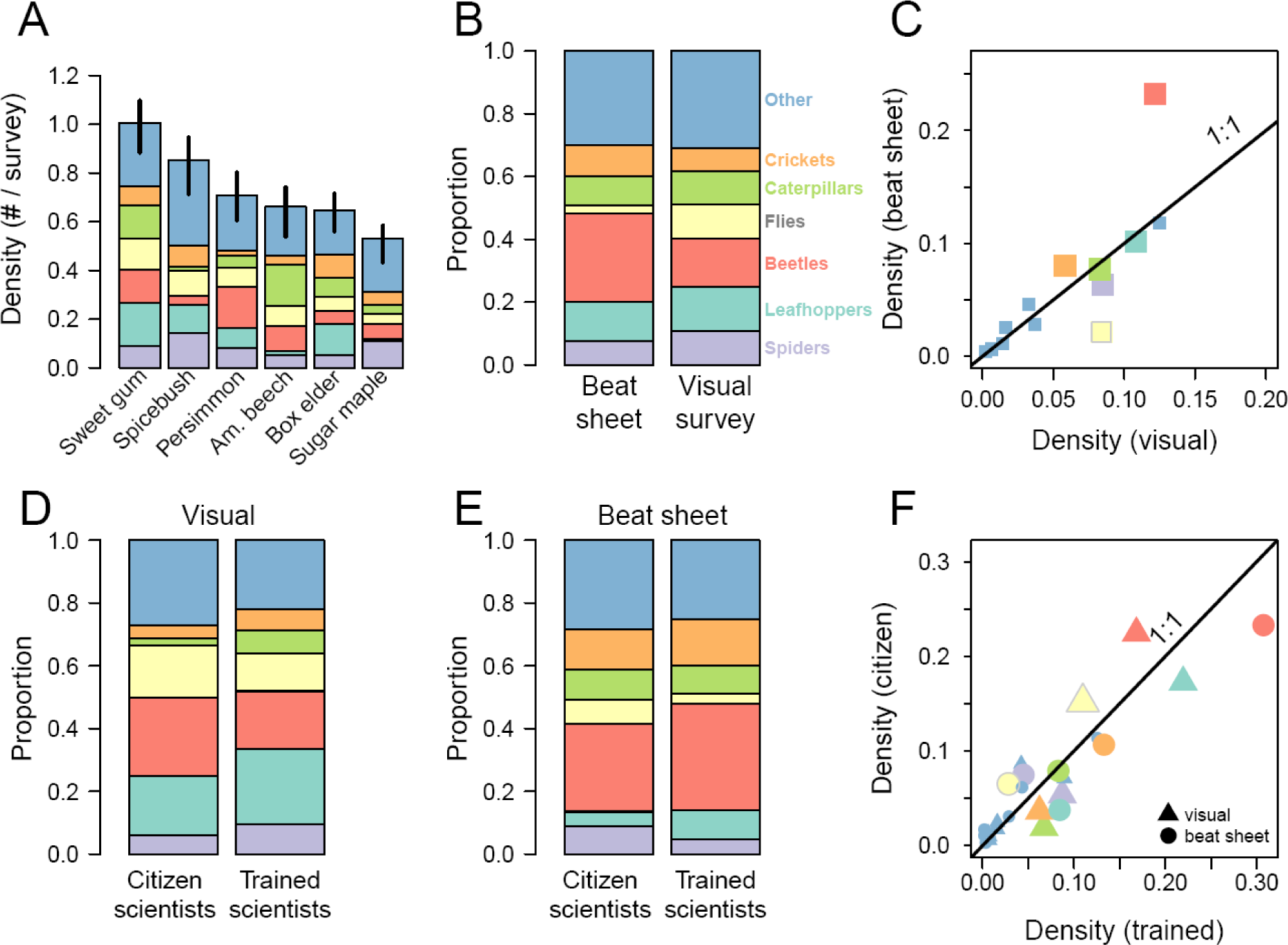
(a) Variation in absolute arthropod density by arthropod group and by tree species for the six most surveyed tree species. Vertical lines indicate 95% confidence intervals on total arthropod density. (b) Variation in the proportion of arthropod groups by survey methodology. (c) Comparison of absolute density estimates of different arthropod groups based on survey methodology. Panels (a-c) include data from both Prairie Ridge and the North Carolina Botanical Garden in both 2015 and 2016. Comparison of the proportion of arthropods observed by citizen scientists versus trained scientists using (d) visual surveys in 2015 and (e) beat sheet surveys in 2016. (f) Comparison of absolute density estimates of different arthropod groups based on whether the data were collected by citizen or trained scientists.

Relative and absolute density estimates for each arthropod group also depended upon survey method (Figure 2b, c; *χ*2 = 284.73, df = 6, *p* < 10^−16^). Beat sheet surveys revealed a greater proportion of Coleoptera (beetles) and a lower proportion of Diptera (flies) compared to visual surveys. A comparison of absolute densities reveals the same discrepancy with respect to the rate at which beetles and flies are observed using the two methods, but also illustrates that density estimates are comparable for most other arthropod groups (r = 0.82, *p* = 0.0004). Notably, caterpillar density estimates were similar using both methods (0.077 versus 0.083 caterpillars/survey for beat sheet and visual surveys, respectively).

Perceived arthropod composition differed between citizen scientist and trained scientist conducted visual surveys (Figure 2d, *χ*^2^ = 44.94, df = 6, *p* < 5e10^−8^). Citizen scientists reported a greater proportion of flies and beetles and a smaller proportion of Auchenorrhyncha (leafhoppers, planthoppers, etc.), Orthoptera (grasshoppers and crickets), and caterpillars compared to the trained scientists, however all differences were within +/- 6%. Absolute density estimates across arthropod taxa were positively correlated between the two groups (Figure 2f, triangles, *r* = 0.84, *p* < 0.0002), and although citizen scientists overestimated fly and beetle density and underestimated caterpillar density relative to trained scientists, these differences were all within 0.05 arthropods/survey.

Using beat sheet surveys, the difference between citizen scientists and trained scientists was less pronounced (Figure 2e, *χ*^2^= 18.34, df = 6, *p* = 0.005), with citizen scientists reporting a slightly greater proportion of Diptera and Araneae and a slightly lower proportion of Auchenorrhyncha and Coleoptera relative to trained scientists. Again, all differences were within +/- 6%. Absolute density estimates were even more strongly correlated across arthropod taxa between the two groups than in the visual survey comparison (Figure 2f, circles, *r* = 0.94, *p* < 0.0001). There was much better congruence in estimates of caterpillar and Orthopteran density in particular using beat sheet surveys compared to visual surveys.

### Phenology

A primary goal of the *Caterpillars Count!* project is to characterize the seasonal fluctuations in arthropods over the spring and summer. The phenology of caterpillars as captured by visual and beat sheet surveys near ground level mirrored the phenology of frass falling from the canopy at PRE (Figure 3a), with less obvious concordance at the NCBG (Figure 3b), although fewer frass data points were available at the latter site.

**Figure 3:**
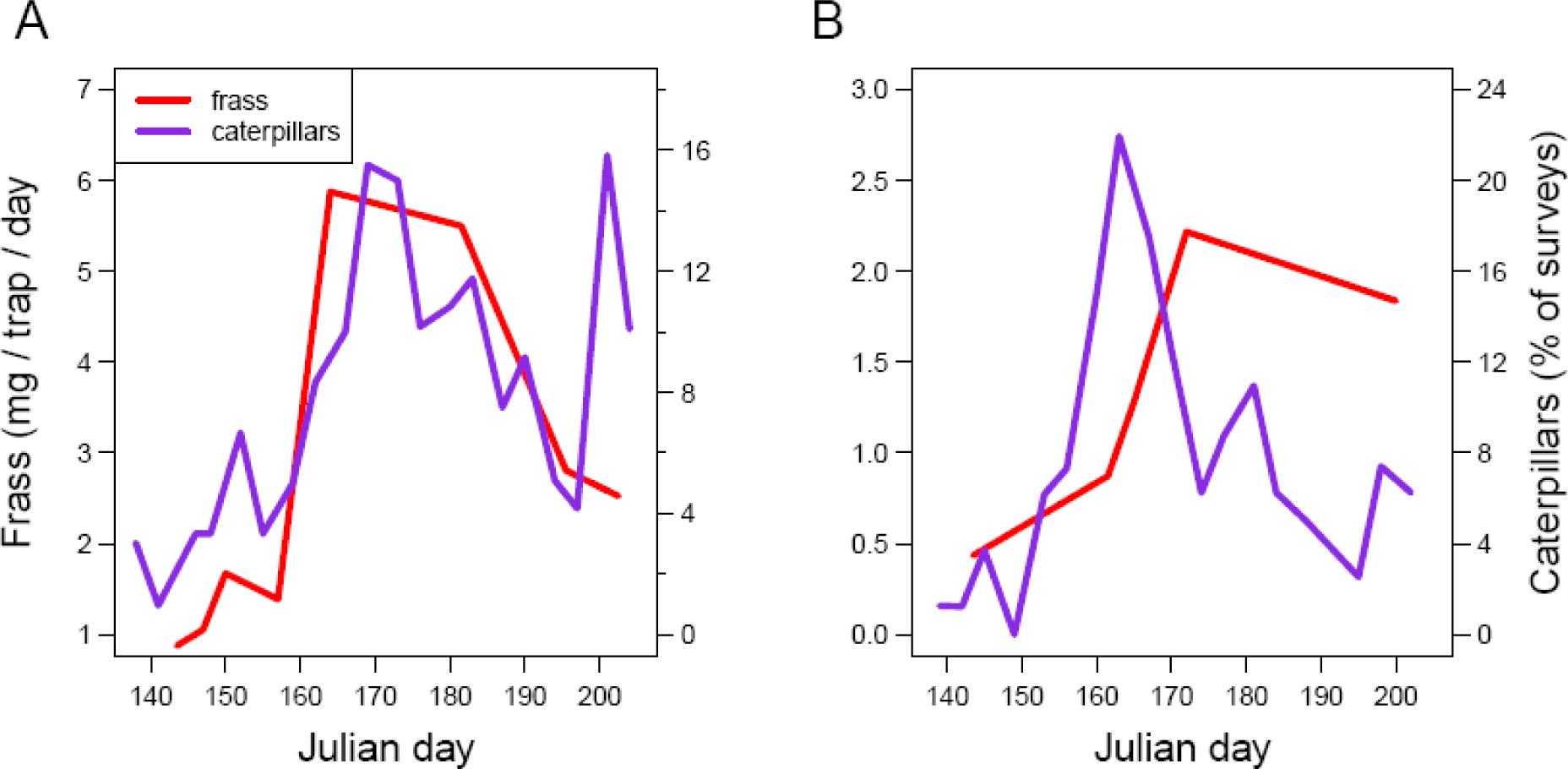
The phenology of caterpillars based on both beat sheet and visual surveys combined (purple) and the phenology of frass collected in frass traps (red) at (A) Prairie Ridge and (B) the NC Botanical Garden in 2015. Frass values were excluded for several dates due to rainstorms.

As expected, arthropods like caterpillars and orthopterans that depend on leaves for food and shelter exhibited low densities in early spring and then increased over the summer (Figure 4a-d, purple lines). Orthopterans continued to increase through mid- to late-July, while caterpillars exhibited a peak in occurrence in mid-June, followed by another in early July. Foliage arthropods in aggregate (caterpillars, orthopterans, beetles, spiders, leafhoppers, and true bugs) exhibit a general positive trend over the dates examined, with less pronounced seasonal peaks due to the more consistent occurrence of some of those other groups like spiders.

**Figure 4:**
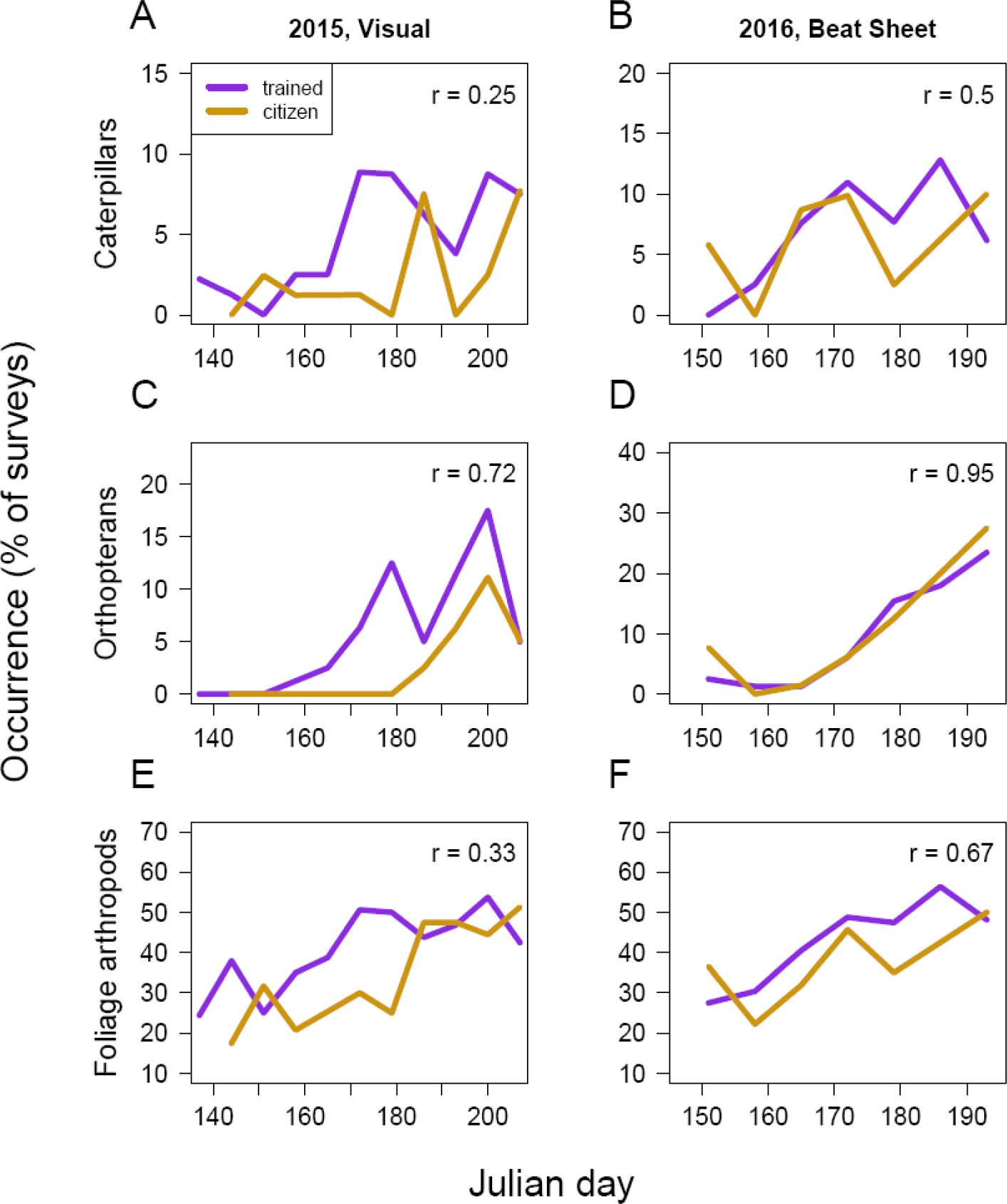
Seasonal phenology in occurrence at Prairie Ridge Ecostation of (a, b) caterpillars, (c, d) orthopterans, and (e, f) a multi-group category including caterpillars, orthopterans, beetles, spiders, leafhoppers, and true bugs based on visual surveys (a, c, e) and beat sheet surveys (b, d, f). Pearson’s correlation coefficient between weekly estimates collected by citizen scientists (orange) and trained scientists (purple) given in the top right.

In 2015 using visual surveys, citizen scientists underestimated the occurrence of foliage arthropods early in the season relative to trained scientists, but estimates converged later in the season (Figure 4a, c, e). Citizen scientists did not observe many caterpillars at all until July. As such, they missed the peak in caterpillar occurrence documented by trained scientists in mid-June, although their observations of a decline in mid-July and subsequent recovery in late July were generally consistent (Figure 4a). Similarly, citizen scientists in 2015 also missed the late June peak in orthopterans but captured the peak in July (Figure 4c).

In 2016, using beat sheet surveys, the phenology recorded by citizen scientists was much more strongly correlated with trained scientist observations (0.50 < *r* < 0.95; Figure 4b, d, f). In particular, citizen scientists identified the same increase and mid-June peak in caterpillar occurrence as trained scientists. Citizen scientists did not actually conduct surveys the week of Julian day 186 when trained scientists identified a second seasonal peak in caterpillars.

### Sampling effort

Estimates of peak caterpillar date were unbiased with respect to the "true" value (Julian day 172) even at low sampling intensity or frequency (Figure 5). However, as expected, 95% confidence intervals around the estimated value were tightest when conducting many surveys at high frequency, or with a low sampling interval. Estimates of peak caterpillar date based on only a small number of surveys or a low frequency of sampling resulted in estimates that were often weeks from the “true” value. In this particular dataset, sampling 30 surveys on a weekly basis led to 95% of estimated peak dates falling within one week of the true date. Increasing the number of surveys conducted per sampling date typically yielded a greater increase in accuracy of the peak date estimate compared to increasing the sampling frequency (Figure 5). For example, doubling the number of weekly surveys from 20 to 40 reduced the confidence interval width by more than 50% (19 days to 9), compared to conducting 20 surveys at double the frequency (19 days to 12).

**Figure 5:**
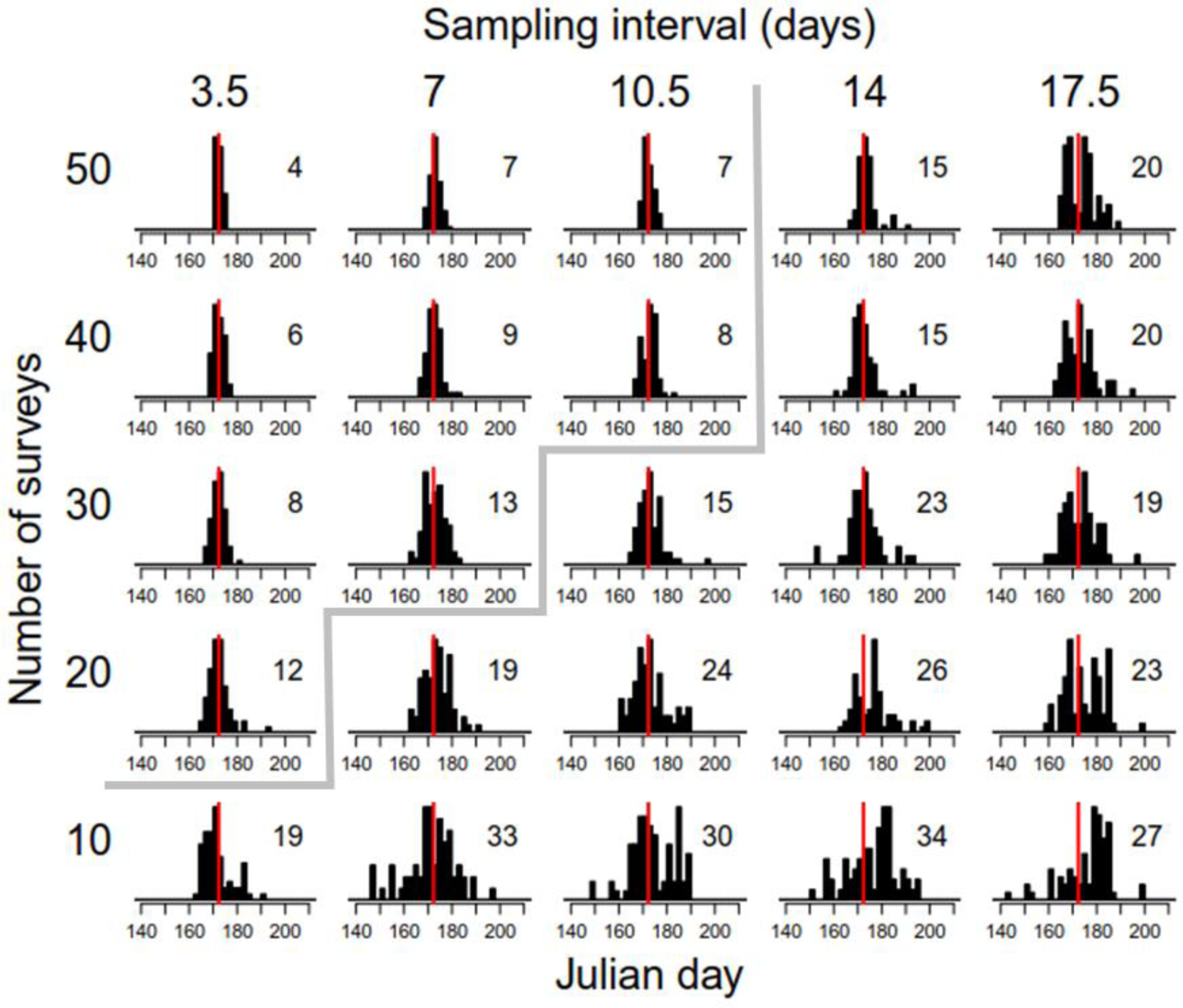
Estimates of peak caterpillar date based on subsampling the Prairie Ridge beat sheet dataset of 2015 to different levels of sampling intensity (rows) and sampling frequency (columns). The “true” estimated peak date based on conducting 60 surveys twice a week was Julian day 172 (June 21; red line). Each histogram indicates the range of peak date estimates based on 60 replicate subsamples for the specified level of sampling frequency and intensity, with the 95% confidence interval width in days in the upper right corner. Sampling combinations above and to the left of the gray line have confidence intervals of 13 days or less.

## Discussion

Foliage arthropod surveys have the potential to shed light on an important and understudied aspect of ecosystem phenology. However, phenology is expected to vary dramatically between regions (Both et al., 2004; Hurlbert and Liang, 2012) and even across local land use gradients (Diamond et al., 2014; White et al., 2002), necessitating the collection of phenology data across broad geographic scales. Here, we have demonstrated the potential of enlisting citizen scientists to collect such data, which could greatly facilitate broad-scale investigations into the wide-ranging impacts of climate change on natural systems. In particular, such data would allow researchers to better interpret the consequences of observed phenological shifts by birds which depend on those arthropod resources (Hurlbert, 2016; Hurlbert and Liang, 2012; Mayor et al., 2017), and shifts by trees and shrubs on which those arthropods depend (Polgar and Primack, 2011; Singer and Parmesan, 2010). These data would also provide a monitoring baseline for assessing arthropod abundance into the future in light of dramatic population declines reported for many groups from across the globe (Dirzo et al., 2014; Hallmann et al., 2017). These preliminary results help inform the best practices for the *Caterpillars Count!* survey scheme that will allow researchers to robustly identify patterns of foliage arthropod density in time and space.

### Phenology at ground level versus the canopy

We found a striking concordance between our ground level survey-based estimates of caterpillar phenology and the canopy level frass-based phenology at Prairie Ridge, suggesting that foliage arthropod surveys conducted near ground level can be used to assess the phenology of higher vegetation strata as well. This correspondence in phenology is consistent with other studies that have found a correlation between lower and upper canopy caterpillar density across trees, years, and season (Cooper, 1988; Holmes and Schultz, 1988). Agreement between caterpillar phenology and frass phenology was weaker at the NC Botanical Garden, with caterpillar density at ground level peaking earlier than frass. In studies where a difference in phenology between strata has been observed, typically it is the canopy that peaks before the understory, when caterpillars migrate down to pupate on the forest floor later in the season (Aikens et al., 2013; Murakami, 2002). The disagreement we observed may instead be due in part to the fact that a large fraction (>70%) of the caterpillars observed at the Botanical Garden occurred in leaf shelters which prevented frass from dropping. In addition, 20% of the survey branches at the Botanical Garden were of the understory shrub spicebush *(Lindera benzoin)* that was not represented at all in the canopy from which frass was being sampled. Monitoring frass phenology at sites where the *Caterpillars Count!* project is implemented will continue to improve our understanding of where and when phenology varies across forest strata, and in some cases could form the basis for a complementary citizen science project.

### Implications for survey methods and sampling scheme

We evaluated two methods for conducting foliage arthropod surveys, visual surveys and beat sheet surveys. In general, the two survey methods yielded very similar results with respect to relative and absolute estimates of arthropod group density based on data collected by trained scientists. As expected, however, each method had its own biases. Flies (Diptera) were underrepresented on beat sheet surveys compared to visual surveys as they tended to fly immediately up and away as soon as a branch was first struck. In contrast, beetles (Coleoptera) were more numerous in beat sheet surveys than visual surveys. Many of the beetles observed in beat sheets were narrow brownish ‘click’ beetles (family Elateridae) which rest flat along twigs. This comparison suggests observers may frequently be overlooking these beetles in visual surveys, although they are quite obvious when lying in a beat sheet. Density estimates for most other groups, including caterpillars, were similar using the two methods. This is interesting given anecdotal observations that some caterpillars, especially those in leaf rolls or sewn between two leaves, are not dislodged by beating, while caterpillars that are extremely cryptic in appearance are more likely to be missed in visual surveys. Although these two groups seemed to be of equivalent abundance such that our two density estimates were comparable, this may not always be the case. Researchers using these data specifically for density estimates will certainly want to take survey method and associated biases into account during analysis, however, phenological metrics of timing which rely on relative, not absolute, indices of abundance should be unbiased. Beat sheet surveys yielded stronger agreement between citizen scientists and trained scientists with respect to density estimates and phenology compared to visual surveys. This was especially true for caterpillars: in 2015 citizen scientists entirely missed the mid-June peak in caterpillar occurrence when conducting visual surveys, while the citizen scientists in 2016 documented patterns similar to the trained scientists using beat sheet surveys. The individual citizen scientist participants differed between 2015 and 2016, indicating that this effect is just as likely to be a participant effect as a survey method effect. Anecdotally, one participant in 2015 was notably less engaged and motivated compared to participants in 2016, highlighting the need to further validate the use of visual surveys in this project. Certainly, not all participants would necessarily have missed the caterpillar peak in 2015. Nevertheless, the ideal methodology is one that is robust to variation in participant ability and motivation. The task of detecting arthropods against a white beat sheet is presumably less subject to error than that of detecting arthropods on an often similarly colored branch, and thus we encourage participants to use beat sheets if possible.

Another advantage of beat sheet surveys in the context of citizen science is the ability to engage and involve younger participants. Although children are not the target participant group for this project, beat sheet surveys require considerably less time and patience than visual surveys, and may be better for youth education programs. Beat sheets are also useful for displaying interesting arthropods to a group, providing an unobstructed view and avoiding the need to have them step up to a branch one at a time. Although constructing a homemade beat sheet is fairly simple and cheap (~$5 in fabric and hardware), it still represents a potential barrier for participants or environmental education centers with limited resources. For that reason alone, we expect that some will choose to conduct visual surveys. Our comparison of the two methods provides an initial suggestion of how to compare data obtained in each, but conducting this methods comparison in other habitats and regions would be useful. Importantly, density estimates of citizen scientists were within 6% of estimates by trained scientists for both survey methods suggesting that either method can yield data useful for addressing research questions.

Finally, we examined how variation in sampling intensity and frequency influenced the perceived date of peak caterpillar occurrence. This is an important question because citizen scientist participants have finite time and resources to dedicate to any particular project, and while estimates of phenology become more precise with increased data collection, the number of participants willing to meet those increased data collection requirements will be smaller (Sauermann and Franzoni, 2015). We found that conducting 30 foliage surveys on a weekly basis provided estimates of peak caterpillar occurrence typically within 1 week of the “true” peak, and recommend this level of effort as a best practice. If a greater sampling effort is possible, increasing the number of surveys conducted per sampling date yields a greater increase in precision compared to investing an equivalent amount of effort in increased sampling frequency and so should be preferred. A smaller number of surveys may still be useful in assessing phenology in a qualitative sense (e.g. determining whether it’s an “early” or “late” year), and we will more rigorously evaluate this possibility as we accumulate more years of survey data.

Because a single foliage survey by an untrained individual conservatively takes about 6 minutes (including sharing observations with others, walking between surveys, etc.), our recommended effort (30 surveys) requires 3 person-hours per week. While some dedicated and interested individuals may participate at this level, they will be in the minority. For this reason, *Caterpillars Count!* will be most easily carried out at centralized locations like environmental education centers that frequently host thousands of visitors each season and have groups of dedicated, regular volunteers eager to contribute toward projects at the site. At centers like these, the data collection effort can be divided up among several people such that, for example, a group of 5 could conduct 30 surveys in less than forty minutes. In this way, individuals interested in participating for only a single day may still contribute to the project within a discrete amount of time and with the assistance of trained and experienced participants. This distributed effort strategy still requires one individual at the site who can coordinate the efforts of other participants, and our experience at Prairie Ridge Ecostation suggests this will require 2 hours per week once the project is up and running.

### Sources of error and bias

Data collection for this project involves three potential sources of error in the context of phenology estimation. First, participants must detect arthropods on survey branches or beat sheets. As discussed above, detectability is expected to be a greater problem for visual surveys due to crypsis, although detectability on beat sheets may still be an issue for arthropods that fly, jump, or run out of the sheet before they can be observed. Nevertheless, for detectability to bias phenological signal, it must vary systematically over time. This may be less of an issue for beat sheet surveys, however, the ability to detect insects on branches via visual surveys almost certainly increases with experience. For sites at which the same individual or individuals conduct visual surveys each week, one might expect observations in the first few survey periods to underestimate arthropod occurrence relative to later in the season. Quantifying exactly how arthropod searching ability improves over time will help determine whether this bias is mostly eliminated after a single day of conducting 5-10 surveys, or if it is likely to persist over a longer period. Nevertheless, to the extent that seasonal arthropods decline in late summer (e.g. July for caterpillars at our study sites), this phenomenon should be well captured by observers regardless of any increases in searching competence.

Second, participants must properly identify arthropods to the appropriate group (Table 1). For groups like caterpillars and spiders, this task will be straightforward. Distinguishing beetles from true bugs and leafhoppers may be more prone to error. We have developed outreach materials including identification keys and cheatsheets to assist participants while they are in the field. We have also developed an arthropod photo identification quiz which is on our website (http://caterpillarscount.unc.edu/quiz/). The quiz may be taken repeatedly with different photos of common foliage arthropods each time, and scores are stored in an internal database by user. Thus, we are able to quantitatively assess the ability of participants to identify the focal taxa. The quiz could thus be used both to filter observations from unreliable users, but also to document any increases in identification ability over time. Finally, when conducting surveys via the mobile app, users may optionally photograph the arthropods they encounter, and these photographs get automatically submitted to the crowdsourcing identification website iNaturalist.org. This feature allows those who are interested to pursue lower taxonomic level identification by experts.

Third, participants must estimate the body length of arthropods to the nearest millimeter. Although much of the US public is less familiar with metric units, having participants calibrate familiar objects like the width of a fingernail or a pencil is fairly straightforward and simple rulers can be drawn on the supports of a beat sheet, or included in the mobile app and arthropod identification guides. Regardless, errors in length estimation will not impact phenology patterns based on occurrence or density. Even in the event that arthropod lengths are used to calculate biomass phenology via length-weight regressions, length estimates are not expected to be biased seasonally in one direction or the other.

### Incentives for participation

Robust survey protocols are necessary but insufficient for ensuring a citizen science project’s success. Equally important are considerations about the motivations and incentives for participating (Hobbs and White, 2012), both from the perspective of potential one-time contributors like weekend visitors to an environmental education center, as well as new potential site coordinators and their regular volunteers. *Caterpillars Count!* provides a context for interested individuals to learn about the natural world around them and to contribute to a broader scientific understanding of arthropod phenology and its consequences in a changing world. The project also provides participants who have affinities to particular *Caterpillars Count!* sites the ability to contribute to something meaningful at that site. We hope the availability of arthropod identification resources, mobile apps for easy data collection, data visualization tools on the project website, and structured learning activities associated with the project will provide additional incentives for environmental educators and others to initiate a *Caterpillars Count!* monitoring scheme.

## Acknowledgements

We would like to thank all of the citizen science participants, especially T. Wilkins and R. Ward, and Hurlbert Lab members that have helped collect data for this project, as well as the environmental educators and others that have provided feedback on protocols, mobile apps, and activities.

